# Assessing the probability and time of weed control failure due to herbicide resistance evolution

**DOI:** 10.1101/2024.04.26.591266

**Authors:** Dana Lauenroth, Chaitanya S. Gokhale

## Abstract

The recurrent exposure to herbicides in agricultural landscapes forces weeds to adapt in a race against extinction. What role newly arising mutations and pre-existing variation play in this evolution of herbicide resistance is critical for developing management strategies. Here, we present a model of rapid adaptation in response to strong selection, capturing complex life cycles of sexual and asexual reproduction and dormancy in a perennial weed. Using a multitype Galton-Watson process, we derive the probability of herbicide resistance evolution and the waiting time distribution until resistant plants appear in the field. We analyse the effect of seed bank dynamics and details of the reproductive system in defining the probability and timing of resistance adaptation in *Sorghum halepense*. Further, we investigate key factors determining the primary source of adaptive variation. We find that even small fitness costs associated with resistance reduce adaptation from standing genetic variation. For herbicide resistance inherited in a (incompletely) dominant fashion, self-pollination also diminishes standing variation for herbicide resistance by increasing the homozygosity. Our study highlights the importance of seed banks for weeds’ adaptive potential, preserving genetic information of forgone selection and prolonging the period in which the population can adapt.

## Introduction

“The assurance of global food security via agricultural yields is […] in need of a highly integrative management approach, in which population geneticists, evolutionary biologists, and weed scientists investigate the nature of resistance adaptation and the factors that affect it to both predict and prevent the evolution of herbicide-resistant weeds.” (Kreiner, Stinchcombe, and Wright, 2017, p. 612) The commercialisation of herbicides during and after the Second World War has vastly improved weed management compared to labour- and cost-intensive mechanical weed control, enabling the intense farming we see nowadays (Oerke, 2006; Tudi et al., 2021). However, analogous to the antibiotic crisis, we ran into a situation where the global presence of resistance towards most known herbicide modes of action threatens the efficacy of chemical weed management (Powles and Yu, 2010; Perotti et al., 2020). The evolution of herbicide resistance is an example of repeated rapid adaptation under human-induced strong selection (Baucom, 2016). It is possible via mutations in genes encoding the targeted protein (target-site resistance) or through nontarget-site alterations like changes in metabolic pathways, reducing the exposure of target proteins to herbicides (Baucom, 2016). Without gene flow from other populations, adaptation can result from standing genetic variation and *de novo* mutations, with existing variation leading to more rapid adaptation in novel environments (Barrett and Schluter, 2008). Identifying the source of adaptive variation is critical to developing strategies to manage herbicide resistance (Kreiner, Stinchcombe, and Wright, 2017). In weeds with substantial standing variation for herbicide resistance, managing the seed bank, for example with stale seedbed preparation, might be vital (Benvenuti et al., 2021). Contrastingly, if de novo mutations serve as the primary source of adaptation, preventing weed seed set by panicle removal or selective herbicide crop topping applications can be a useful (Tidemann et al., 2021).

Standing variation for herbicide resistance was present in populations of weeds such as annual ryegrass (*Lolium rigidum*) and blackgrass (*Alopecurus myosuroides*) that were never exposed to herbicides, even predating their introduction in blackgrass (Preston and Powles, 2002; Délye, Deulvot, and Chauvel, 2013). Recent genomic analysis and simulations by Kersten et al. (2023) highlight the prevalent role of standing genetic variation in target-site resistance adaptation of the annual weed blackgrass. Conversely, in species like common waterhemp (*Amaranthus tuberculatus*) and foxtail millet (*Setaria italica*), high resistance costs might diminish standing variation, making novel mutations a more likely source of adaptation (Vigueira, Olsen, and Caicedo, 2013). Genomic data of several waterhemp populations supports a prevailing role of *de novo* mutations in target-site resistance of this species (Kreiner et al., 2022). Yet, the precise determinants of the primary source of adaptive variation in herbicide resistance remain unclear.

Branching processes are well-established in biological studies for modelling population evolution (Harris, 1963; Athreya and Ney, 1972; Kimmel and Axelrod, 2002; Santer and Uecker, 2020). They have been used to model populations at risk of extinction due to low reproduction rates after an environmental change, investigating how the emergence of an adapted mutant could prevent population extinction via evolutionary rescue (Orr and Unckless, 2008; Uecker, Otto, and Hermisson, 2014; Azevedo and Olofsson, 2021). Multitype branching processes can yield further information about the waiting time until such a mutant appears (Serra and Haccou, 2007; Alexander, 2016). Despite recognition of the value of concepts from evolutionary rescue for understanding population genetics of herbicide resistance evolution (Neve et al., 2014; Kreiner, Stinchcombe, and Wright, 2017), this approach is lacking in weed science literature. Our work addresses this gap.

Using multitype Galton-Watson processes, we analyse the probability and timing of herbicide resistance evolution in perennial weeds causing treatment failure. In this context, our study focuses on the influence of specific weed traits—asexual propagation, self-pollination and seed dormancy. We emphasise the role of the seed bank in the evolution of herbicide resistance, preserving genetic information of forgone selection and prolonging the time for adaptation. Further, we study critical ecological and evolutionary factors that modify the role of *de novo* mutations and standing genetic variation in the evolution of herbicide resistance. The seed bank size determines the absolute frequency of pre-existing variants for target-site resistance, and the seed production capacity of the weed defines the number of newly arising mutations, affecting their relative contribution to resistance adaptation. Self-pollination and fitness costs associated with resistance both reduce adaptation from the standing genetic variation. Our theoretical results explain the variation in primary sources of adaptive variation observed in different weed species, determining effective strategies for herbicide resistance management.

## Methods

We explore the dynamics of rare beneficial alleles within large populations facing sudden environmental shifts, like herbicide application. In particular, we model the appearance and spread of target-site mutations, conferring resistance to the herbicide, in a primarily sensitive weed population declining in response to the treatment.

We illustrate our modelling framework on the example of a perennial weed, *Sorghum halepense* (Johnsongrass), which reproduces sexually via seeds and asexually through rhizomes (Warwick and Black, 1983; Peerzada et al., 2023). The weed enters dormancy during winter, surviving as seeds and rhizomes underground until new young plants emerge in spring and mature over the season to produce new rhizomes and seeds (Warwick and Black, 1983; Peerzada et al., 2023). These seeds, known for their high dormancy, contribute to forming a seed bank, ensuring the species’ viability in the soil (Warwick and Black, 1983; Peerzada et al., 2023). We illustrate this cyclical life pattern of Johnsongrass along with the further plant stages and herbicide application in ***Figure 1*** a. The mathematical representation of the model is shown in ***Figure 1*** b.

**Figure 1.**
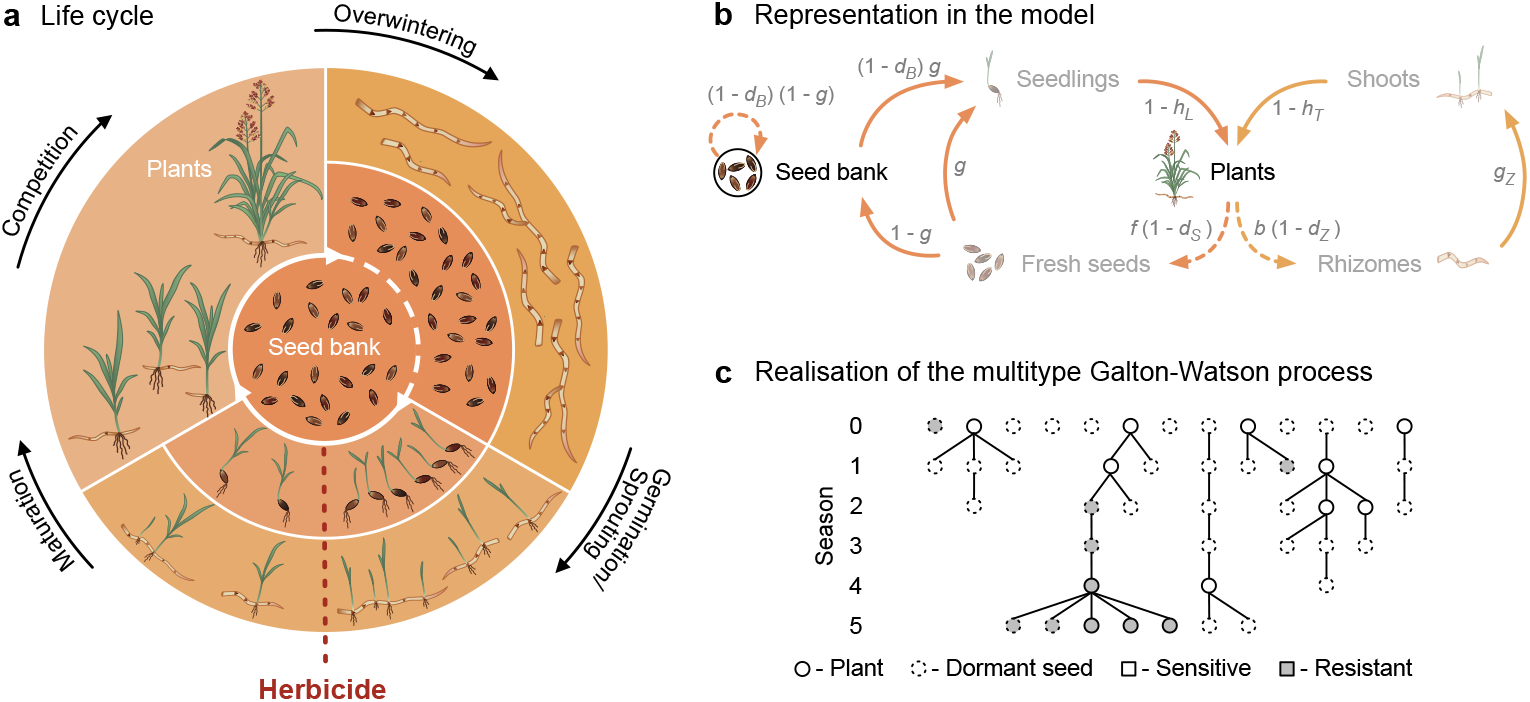
Schematic illustration of Johnsongrass’ life cycle and its representation in our model. a, Diagram of Johnsongrass’ life cycle. Johnsongrass reproduces sexually via seeds (inner ring) and asexually through rhizomes (outer ring). Seeds can stay dormant in the ground for several years, forming a seed bank (central circle). New seeds and seeds from the seed bank might germinate in spring or stay dormant as part of the seed bank (expressed by the dotted line). Rhizomes give rise to shoots in the first spring after their production. Herbicide application (red dotted line) can kill susceptible seedlings and shoots. Surviving plants compete for resources as they mature. The aboveground plant material dies in winter, and Johnsongrass overwinters as seeds and rhizomes in the ground (adapted from Lauenroth and Gokhale (2023)). **b, Representation of the life cycle in our model**. The left part of (b) corresponds to the sexual reproduction of Johnsongrass, and the right part represents the asexual propagation. Our model tracks the seed bank and reproducing plants; the other life history stages depicted transparently are not explicitly modelled. Solid arrows depict within-season dynamics, and dashed arrows show dynamics between seasons. Survival probabilities and fecundity are shown in grey next to the corresponding arrows: number of seeds *f* and buds on the rhizome *b*; winter survival of fresh seeds 1 − *d*_*S*_, seeds in the seed bank 1 − *d*_*B*_ and rhizome buds 1 − *d*_*Z*_ ; probability of seed germination *g* and rhizome bud sprouting *g*_*Z*_ ; herbicide survival of seedlings 1 − *h*_*L*_ and shoots 1 − *h*_*T*_. **c, Illustration of a realisation of our multitype Galton-Watson process**. Shown is the phylogeny of a branching process resembling a herbicide-treated weed population over six seasons. Open circles depict plants, and dashed circles represent dormant seeds in the seed bank; sensitive individuals are white, and grey are individuals resistant to the herbicide.

In Lauenroth and Gokhale (2023), we presented a deterministic model of Johnsongrass’ life cycle capturing all life-history stages and intraspecific competition between the plants, allowing the prediction of long-term population dynamics (***Figure 1*** a). In the present study, we are interested in the probability and timing of herbicide resistance evolution. With herbicide exposure, the common sensitive allele, carried homozygously by the majority of the population, becomes maladaptive. Target-site mutations, in particular, can endow a high level of resistance (Scarabel et al., 2014), warranting that the first resistant plant surviving till reproduction almost certainly ensures population survival. We, therefore, concentrate on adult plants and seeds within the seed bank, counted at the end of each season before the newly produced seeds enter the seed bank. Focussing on the invasion dynamics, our branching process model captures the early phase of adaptation in which the resistance allele is rare.

### Branching process model

We consider a total of *m* genotypes, which may correspond to different mutations at one locus or several loci or take different levels of ploidy into account, denoted by {1, 2, …, *m*}, where genotype 1 is the prevailing sensitive type, and the rest are rare. This setup leads to tracking 2*m* types {1, 2, …, 2*m*}, with the first *m* being plants and the latter half representing seed genotypes. We define *C ⊂* {1, 2, …, *m*} as the set of plant types expected to grow positively, termed supercritical types, while the remainder, {1, 2, …, *m*}∖*C*, are subcritical, facing extinction unless a successful mutant arises. Producing at most one offspring, seed types {*m* + 1, *m* + 2, …, 2*m*} are as well subcritical.

We formally assume all individuals—seeds and plants—perish at the end of the season and produce offspring independently, constituting the following season’s population. A seed that stays dormant and part of the seed bank thus has one identical offspring—itself—in the next season. While seeds have at most one offspring in the next season, plants can generate multiple offspring, either as new plants or seeds (illustrated in ***Figure 1*** c). The population in season *n* is represented by the random vector ***Z***_*n*_ = (*Z*_*n*, 1_, *Z*_*n*, 2_, …, *Z*_*n*, 2*m*_), where the first *m* entries count the different plant types and the second half contains absolute genotype frequencies in the seed bank. The offspring of type *i* individuals are described by independent and identical distributed random vectors 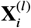, taking values in 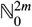, where the entries 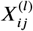 give the number of type *j* offspring produced by the *l*-th type *i* individual. The population vector ***Z***_*n*_ is then for *n* ∈ ℕ_0_ recursively defined by a given initial population 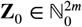 and

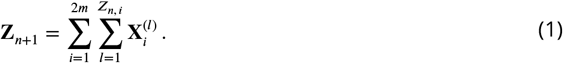

The time discrete homogeneous Markov chain 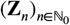 with state space 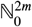 is a multitype Galton-Watson process modelling the weed population dynamics over time. We show a realisation of a simplified process in ***Figure 1*** c.

#### Offspring generating function

We assume plants produce a Poisson-distributed number of seeds and rhizome buds, which might die over winter before they potentially develop into plants. A herbicide application can kill sensitive plants, while dormant seeds are unaffected. The processes, including survival and development, are considered independent among units, leading to offspring numbers that follow independent Poisson distributions (Acharya et al., 2014). We assume vegetative propagation produces genetically identical offspring, neglecting somatic mutations. Seeds can differ genetically from the parent plants due to Mendelian inheritance and mutations—mutations and back mutations at a given rate μ. Johnsongrass primarily self-pollinates, but since cross-pollination is possible at sufficiently high densities (Warwick and Black, 1983), our model captures a continuum, presuming abundant pollen results in complete ovule fertilisation. Our focus is on the proliferation of rare alleles, positing that a prevalent homozygous genotype, labelled genotype 1, predominates while alternative allele carriers are rare. We simplify by assuming these rare types do not engage in cross-pollination with each other, but type 1 plants can cross-pollinate with all genotypes. Let *χ*_*S*_ and *χ*_*R*_ denote the absolute frequency of the sensitive type 1 and rare resistant plants, respectively. The fraction of sensitive plants pollinated by resistant plants’ pollen is 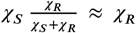 and the fraction cross-pollinated by pollen from sensitive plants is 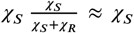. It is, hence, natural to count all offspring produced by matings between type 1 and rare type plants as the rare type’s offspring. Since the inheritance does not depend on which mating partner contributes the male or female gametes, this altered offspring assignment does not constitute a deviation from Mendelian inheritance. These approximations of the cross-pollination allow us to capture sexual reproduction in the asexual Galton-Watson process. Taken together, the genotype *j* plants and dormant seeds produced by a plant of genotype *i* are independent and Poisson distributed, with means *λ*_*ij*_ and *λ*_*i m*+*j*_ respectively, with Johnsongrass-specific parameters detailed in the Supplementary Information.

Seeds in the seed bank face three possible outcomes: germination into a plant of the same genotype *i* with probability *α*_*i*_, remaining dormant with probability *β*, or disappearing without offspring with probability *γ*_*i*_.

This defines the offspring distributions 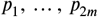 of the branching process, where *p*_*j*_ (***r***) specifies the probability of a type *j* individual leaving offspring vector ***r*** in the following season. For plants of genotype *i*, the generating function of their offspring is then given by

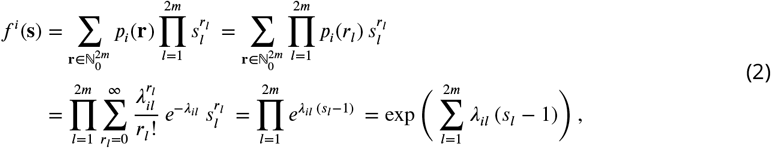

where the second equality holds because we have independent Poisson distributions resulting from Poisson sampling (Acharya et al., 2014). The order of summation can be changed to reach the third equality due to the absolute convergence of the series on [−1, 1]. For seeds of genotype *i*,

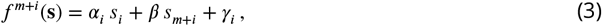

is the offspring-generating function. The offspring generating functions for the different genotypes of plants and seeds will be combined in the vector ***f*** (***s***) := *f* ^1^(***s***), *f* ^2^(***s***), …, *f* ^2*m*^(***s***) .

#### Extinction probabilities

A branching process ultimately either becomes extinct or grows without bounds. The interest lies in the extinction and survival probabilities, which reflect the chance of treatment success (weed eradication) or failure (weed proliferation). We denote the probability that a process started by one type *i* individual becomes extinct with *q*_*i*_. We summarise the type-specific extinction probabilities in a vector ***q*** = (*q*_1_, *q*_2_, …, *q*_2*m*_). The extinction probability vector ***q*** is the solution of

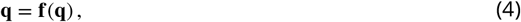

in the cube [0, 1]^2*m*^ closest to **0** (Sewastjanov, 1974, p.115). The vector **1** invariably satisfies Equation (4), indicating that in the absence of any other solutions within [0, 1]^2*m*^, the associated branching process is sure to become extinct, with a probability of 1, regardless of the initial population.

The first *m* equations in (4) stem from the offspring-distributions of the plants, yielding extinction probabilities of processes initialised by a plant; the second half corresponds to processes starting with a single seed. Rearranging the bottom half of the equations in (4),

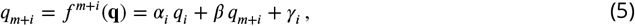

we can express the extinction probabilities of processes starting with a single seed in terms of the extinction probabilities when starting with a plant of the same genotype,

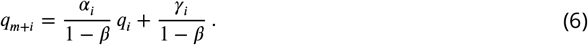

Substituting this relation into the top half of the equations in (4),

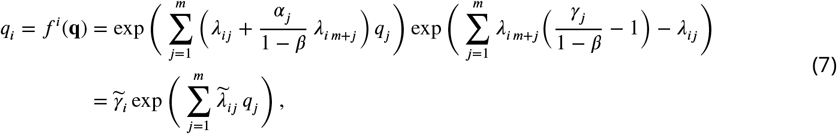

where 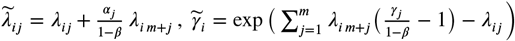, *i, j* ∈ {1, 2, …, *m*}, we can effectively halve the dimension of the equation system.

#### Probability of treatment failure

The extinction probabilities, ***q***, for processes starting with one individual enable us to determine the probability of extinction or survival for a process initiated with any initial population **z**_0_, corresponding to weed treatment success or failure.

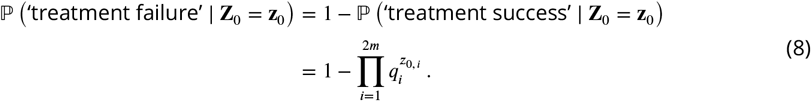

The initial population distribution influences the extinction probability. Assuming Poisson distributed plant and seed numbers, with means *n*_*P*_ and *n*_*B*_, the initial counts of each type are independent and Poisson distributed, with means *n*_*P*_ *v*_*i*_ and *n*_*B*_ *v*_*i*_, for seeds and plants respectively, where *v*_*i*_ denotes the frequency of genotype *i* in the pre-existing variation (Acharya et al., 2014). The probability of treatment failure is then analogue to (2) given by,

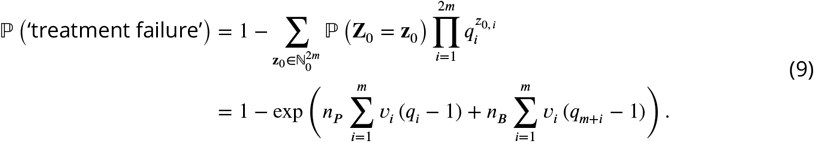

We can use Equation (9) to analyse the role standing genetic variation and *de novo* mutations play in herbicide resistance adaptation. By setting *v*_1_ = 1 and *v*_*i*_ = 0 for *i* > 1, corresponding to the start population consisting of only sensitive homozygotes, we get the probability of treatment failure through *de novo* mutations

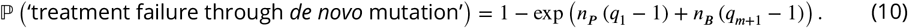

Excluding mutations, we get the offspring generating function without mutation 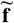 and corresponding mutation free extinction probabilities as 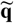. To calculate the probability of treatment failure through standing genetic variation, we insert 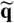 in Equation (9),

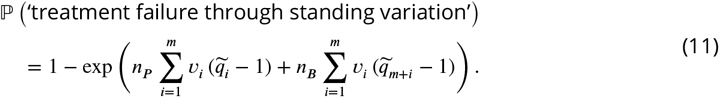

Adaptation through *de novo* mutations and pre-existing genetic variation are intertwined, potentially contributing to population survival. Unlike the approach of Hermisson and Pennings (2005) for adaptation from pre-existing variation and new mutations, our model does not separate the total adaptation probability into contributions from standing variation and *de novo* mutations as we have also included back mutations. Thus, Eqs. (10) and (11) represent the adaptation probabilities in populations without pre-existing target-site resistance or those that cannot mutate, relying solely on existing variation, respectively.

#### Waiting times

The time until resistant plants establish, allowing for the survival of the weed population against herbicide control, is critical. We apply findings from Alexander (2016) to calculate the probability that resistant plant types have not appeared and to determine the distribution of waiting time for their emergence.

*C* denotes the set of supercritical types corresponding to resistant plants. We calculate the probability of *C* not appearing by a specific time. A particular type has yet to emerge by generation *n* if it did not appear by generation *n* − 1 and no generation *n*− 1 individual produced such offspring. Therefore, the probability Π_*i*_(*n*) that resistant plants have not appeared in a type *i* individual’s lineage by generation *n* is given by

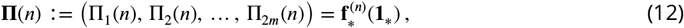

where 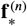 denotes the *n*-th iteration of the offspring generating function ***f*** with the components corresponding to the resistant plants *C* fixed at zero, i.e. 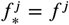 unless *j* ∈ *C* when it is 0. **1**_*\**_ is the vector of length 2*m* where all entries are 1 except for the entries of the resistant plants *C*, which are again 0.

The random variable *N* denotes the time until the first appearance of a resistant plant (first resistant plant establishes and sets seed). The probability of a resistant plant appearing until a given generation *n* is the complementary probability of no such plant appearing until generation *n*. Hence, we can derive the cumulative distribution of *N* from the probabilities Π_*i*_(*n*) in (12) that no resistant plant has appeared in the lineage of a type *i* individual by generation *n*. If the process starts with just one individual of type *i*,

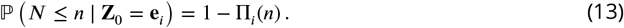

No resistant plants arise in the branching process until some time, if and only if no such individuals appear in any of the independent lines. Thus, we can express the cumulative function of *N* analogue to the treatment failure probability in (9) as

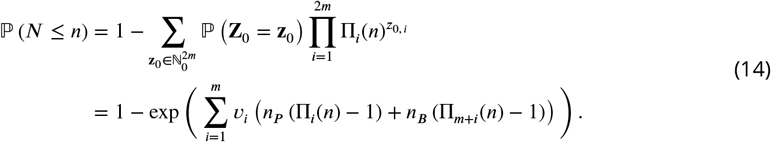

Since the process is time discrete, the probability mass function of the waiting time *N* until the appearance of a resistant plant is given by

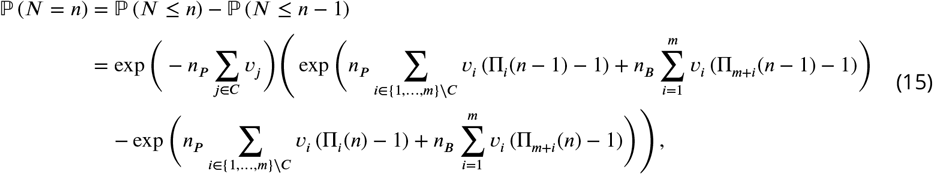

for *n* ∈ ℕ and *ℙ* (*N* = 0) = 1 − exp (−*n*_*P*_ Σ_*j*∈*C*_ *v*_*j*_). Since the population might go extinct without any resistant plant appearing, *ℙ* (*N* = ∞) > 0. Even if a resistant plant exists at some time in the population, it might still face extinction, 0 < *ℙ* (*N* = ∞) < *ℙ* (‘treatment success’).

In the following, we assume a diploid genetic architecture with one allele conferring herbicide resistance at a single biallelic locus (Lauenroth and Gokhale, 2023). We consider the case of incomplete dominance of resistance, with homozygous resistant plants not affected by the herbicide and heterozygous plants being killed with an efficacy half as high as for sensitive homozygotes. We implement an incompletely dominant fitness cost *c* on seed production associated with resistance, such that heterozygotes suffer half the reduction in seed production capacity as plants carrying the resistance allele homozygously.

### Computer simulations

We simulated our branching process model to observe the dynamics of reproducing plants and the seed bank in a Johnsongrass population (implemented in MATLAB version 9.14.0 (R2023a)). Initial populations of plants and seeds were based on independent Poisson distributions, using the product of expected gene frequencies in untreated populations *v*_*i*_ and the number of plants *n*_*P*_ or seeds *n*_*B*_ as means. We estimated the standing variation for target-site resistance ***v*** = (*v*_1_, *v*_2_, …, *v*_*m*_) using a deterministic density-dependent model (Lauenroth and Gokhale, 2023), simulating over 1000 years without management to achieve mutation-selection balance (***Figure S1*** a). The plant and seed offspring of type *i* plants followed independent Poisson distributions with means 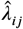 and 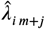, respectively, allowing for cross-pollination between all possible genotypes. Simulations included density-dependent reproduction effects, modifying fecundity based on high population density as per Eqs. (11) and (12) in Lauenroth and Gokhale (2023). The genotype *i* seeds had multinomially distributed plant and seed offspring of the same genotype—seeds germinating and remaining dormant. The process was deemed extinct if no plants or seeds remained, and treatment was considered failed once resistant plants were present (yielding a weed extinction probability of less than 10äunder the standard parameter set in ***Table S1***). We recorded the proportions of extinctions and treatment failures, extinction times and time of appearance of the first resistant plant. The code is available on GitHub. We provide separate folders for generating all figures in the manuscript with subfolders *Simulations* containing the Matlab files Dynamics.m that generates the population dynamics where relevant.

## Results

The survival of a weed population under herbicide treatment is tied to the establishment of resistant plants on the field. For our standard parameter set given in ***Table S1***, lineages of sensitive plants and seeds face extinction with a probability close to one. With at most one successor in the next season, heterozygous and homozygous resistant seeds also have high extinction probabilities of around 0.88 and 0.75. Plants tracked by our model are the successful plants that reproduce. The high level of resistance endowed by the target-site mutation and the high fecundity of Johnsongrass cause the extinction probabilities of lineages started by a resistant plant to be close to zero, with 8.6 · 10^−66^ for heterozygotes and 1.3 · 10^−109^ for homozygotes. Hence, once a resistant plant appears on the field and survives until reproduction, its lineage is extremely unlikely to go extinct (***Figure S3*** a).

### Source of adaptive variation

#### Considering self-pollination

Herbicide resistance in weeds can arise from standing genetic variation and *de novo* mutations, determining the probability and rate of adaptation. The source of adaptative variation is thus relevant for the success of management strategies. We define the establishment of resistant plants under herbicide leading to population regrowth as treatment failure, corresponding to the evolutionary rescue of a population through successful mutants (Gomulkiewicz and Holt, 1995; Neve et al., 2014; Bell, 2017; Kreiner, Stinchcombe, and Wright, 2017). By calculating probabilities of treatment failure through standing variants (Eq. (11)) or *de novo* mutations (Eq. (10)), we assess their impact on adaptation. A fitness cost *c* associated with resistance considerably lowers the probability of adaptation from standing variants by decreasing resistant plants’ relative fitness in the absence of herbicides, lowering their abundance (***Figure 2*** a) (see Lauenroth and Gokhale (2023) for more details). The probability of treatment failure, influenced by field size, follows a sigmoidal curve, with larger fields or higher initial densities (effectively larger population sizes) increasing treatment failure probabilities (***Figure 2*** b). Self-pollination reduces adaptation from standing genetic variation caused by the increased homozygosity, decreasing the frequency of resistant individuals. However, the higher number of resistant homozygotes in the standing variation of self-pollinated weeds is advantageous if the mutation is highly recessive. In contrast, we expect the same number of de novo mutations regardless of the type of pollination (see Appendix 1 and ***Figure S4*** for more detail), causing treatment failure with similar probabilities if heterozygotes have high fitness. If heterozygous plants can not restore positive population growth, selfing again facilitates adaptation by efficiently generating resistant homozygotes. For self-pollinating species like Johnsongrass, considerable resistance costs make *de novo* mutations a likely source of target-site resistance.

**Figure 2.**
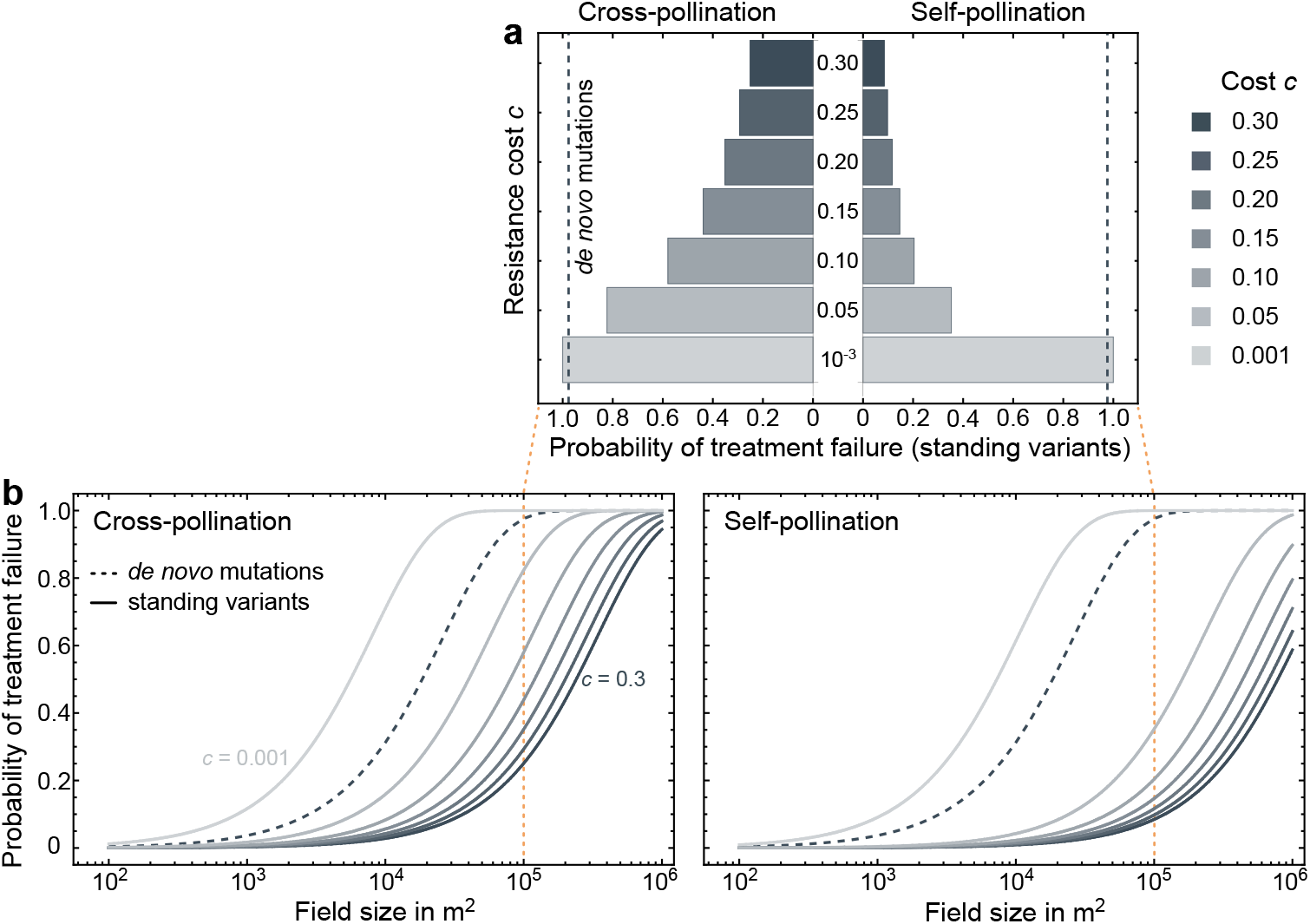
Importance of *de novo* mutations and standing genetic variation for herbicide resistance adaptation depending on the associated resistance cost and self-pollination. We show the probabilities of treatment failure through standing genetic variants derived from Equation (11) under fitness costs on seed production between 0.1 and 30 %. Darker shades indicate higher costs of resistance. The dashed lines show the probability of treatment failure due to *de novo* mutations obtained from Equation (10). The left side reflects a fully cross-pollinating population, and the right represents a self-pollinated population. **a, Probabilities of treatment failure in a field of** 10^5^ **square metres. b, Treatment failure probabilities as a function of field size**. The figures use the default parameter set as defined in ***Table S1*** except when specified otherwise.

#### Considering seed bank density and fecundity

The seed bank acts as a reservoir for genetic variation, with a larger seed bank increasing the chances of resistant plant emergence and treatment failure (***Figure 3***). The probability of new resistant mutations is tied to plant fecundity; thus, any decrease in seed production substantially lowers the probability of treatment failure through *de novo* mutations (lines in ***Figure 3***). Besides affecting the overall probability of control failure, the initial seed bank size and fecundity considerably impact the relative importance of standing variation and *de novo* mutations as a source of adaptation, highlighting the necessity of considering the weeds ecology and infestation history in the design of resistance management.

**Figure 3.**
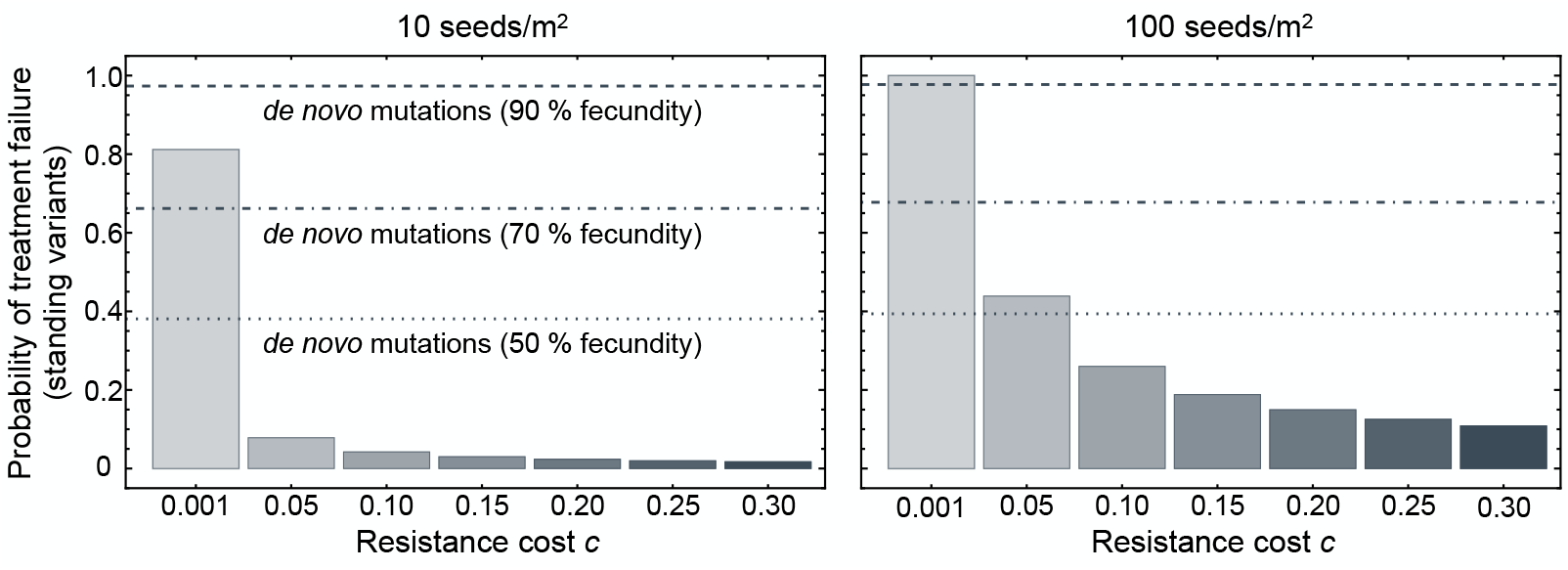
Role of seed bank density and fecundity for the relevance of *de novo* mutations and standing genetic variation in herbicide resistance evolution. We show the probabilities of treatment failure through standing genetic variants derived from Equation (11) under fitness costs on seed production between 0.1 and 30 %. Darker shades indicate higher costs of resistance. The lines show the probability of treatment failure due to *de novo* mutations obtained from Equation (10) for 90 % of maximum fecundity (dashed), 70 % of maximum fecundity (dashed-dotted) and 50 % of maximum fecundity (dotted). The left panel corresponds to an initial seed bank density of 10 seeds per square metre, and the right panel reflects 100 seeds per square metre. The figures use the default parameter set as defined in ***Table S1*** except when specified otherwise.

### Probability and time of control failure

#### Considering asexual propagation

The control of rhizomes is only partial as the herbicide active ingredients are not fully translocated into the rhizomes (Beasley, 1970; Tuesca et al., 1999). Potential resprouting from surviving rhizomes reduces the herbicide efficacy on rhizome shoots compared to seedlings (Vidrine, 1989). We varied the weed’s investment in sexual and asexual reproduction to analyse vegetative proliferation as a strategy to overcome control. Specifically, we fixed the total expected number of direct plant offspring one plant has in the following seasons without control measures and varied the proportion of asexually generated offspring, implicitly assuming that surviving sexual and vegetative offspring are equally costly. Though the mutational influx is less, since we do not consider mutations in vegetative growth, plants that invest more in asexual reproduction are more likely to escape control (***Figure 4*** a). Vegetative proliferation potentially improves population survival under herbicide treatment if standing variation for target-site resistance exists since the offspring of herbicide-resistant individuals are themselves resistant. Not all sexually generated offspring will be resistant, especially for heterozygous mother plants or due to cross-pollination in predominantly susceptible populations. However, clonal reproduction per se is not beneficial here (dashed line in ***Figure 4*** a), likely due to the omission of somatic mutations and the high fitness of heterozygotes. Nevertheless, rhizomes as an organ of vegetative reproduction increase the persistence as a result of reduced control by the herbicide, buying the weed population time to generate a successful mutant (***Figure 4*** b). Thus, we expect control failure by herbicides to be likelier and to occur later in weeds investing more in asexual propagation (***Figure 4***).

**Figure 4.**
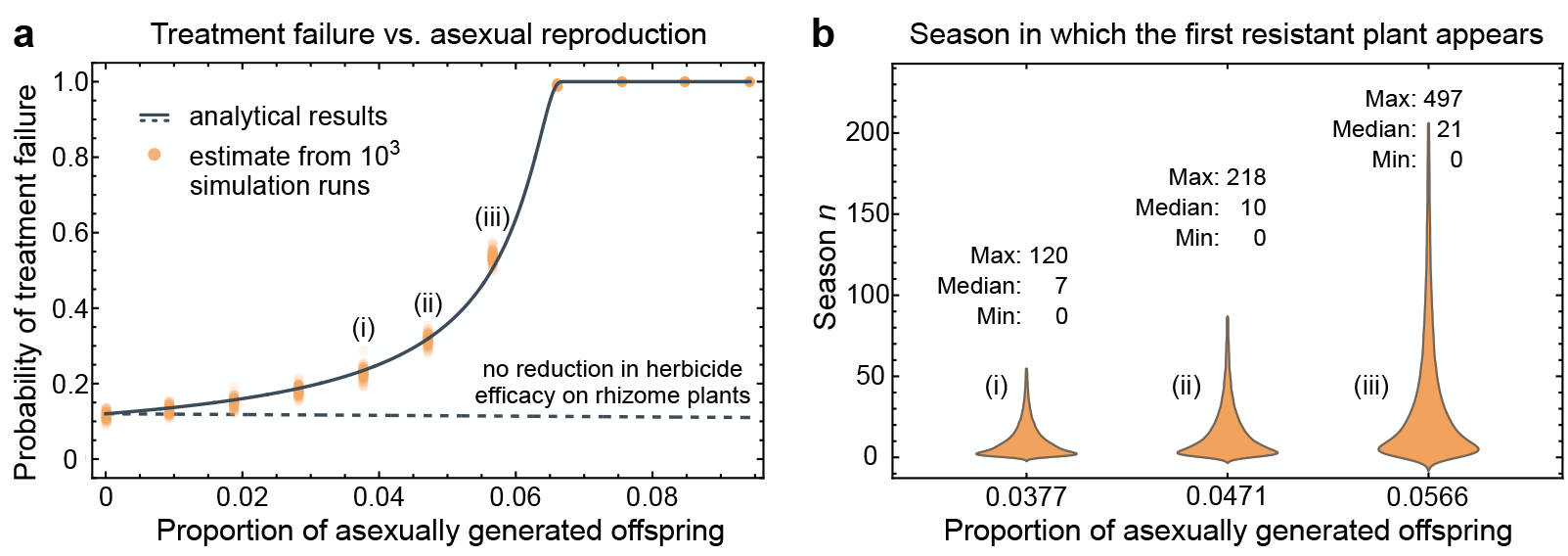
Impact of asexual propagation on herbicide resistance adaptation. We show analytical results and simulation estimates for varying proportions of asexually generated offspring under a fixed expectation of plant offspring one plant generates without control measures. **a, Variation of the treatment failure probability with the proportion of asexually generated offspring**. The grey lines show the probability of treatment failure derived analytically from Equation (9); as a solid line under reduced herbicide efficacy on rhizome shoots (see ***Table S1***) and as a dashed line for the herbicide efficacy on rhizome shoots equal to the efficacy on seedlings (*h*_*T*_ = *h*_*L*_). The closed orange circles are estimates from 10^3^ simulation runs with 10^2^ replicates each. **b, Distribution of time until the appearance of the first resistant plant for different proportions of asexually generated offspring**. The violin plots show smoothed probability density estimates of the season where the first resistant plant is established on the field. The density estimates were generated from 10^5^ simulation runs for the proportions of asexually generated offspring marked in (a). The maximum, median and minimum are displayed on the violin plots. The figures use the default parameter set as defined in ***Table S1*** except when specified otherwise.

#### Considering the seed bank

A seed bank stores genetic information from several past generations, preserving adaptation to previous herbicide applications. Further, seed banks buffer changes in population size induced by the herbicide. How long seeds stay in the seed bank and how many of them ultimately germinate is determined by the germination probability *g* and the seed mortality *d*_*Z*_. High germination leads to a fast turnover—weak seed bank—and low germination causes a slow turnover—strong seed bank. To analyse the effect of seed bank strength, we varied the germination probability of weed seeds and the seed bank mortality while maintaining the total number of sexually generated offspring per plant constant. We simulated an earlier treatment with the herbicide followed by ten seasons without control measures, yielding the composition of the start population (***Figure S1*** b). Populations with a stronger seed bank have a higher probability of escaping from following herbicide applications (***Figure 5*** a) with higher probabilities of resistant plants establishing early on (***Figure 5*** b) since the change in allele frequencies induced by the previous application is preserved in the seed bank. Weaker seed banks with higher germination, in turn, slow the population decline after treatment onset, enabling the later appearance of herbicide-resistant plants due to *de novo* mutations (***Figure 5*** b). After several seasons of herbicide application, the weaker seed bank is depleted, and resistant plants are again slightly more likely to emerge in populations with seeds still preserved in a strong seed bank.

**Figure 5.**
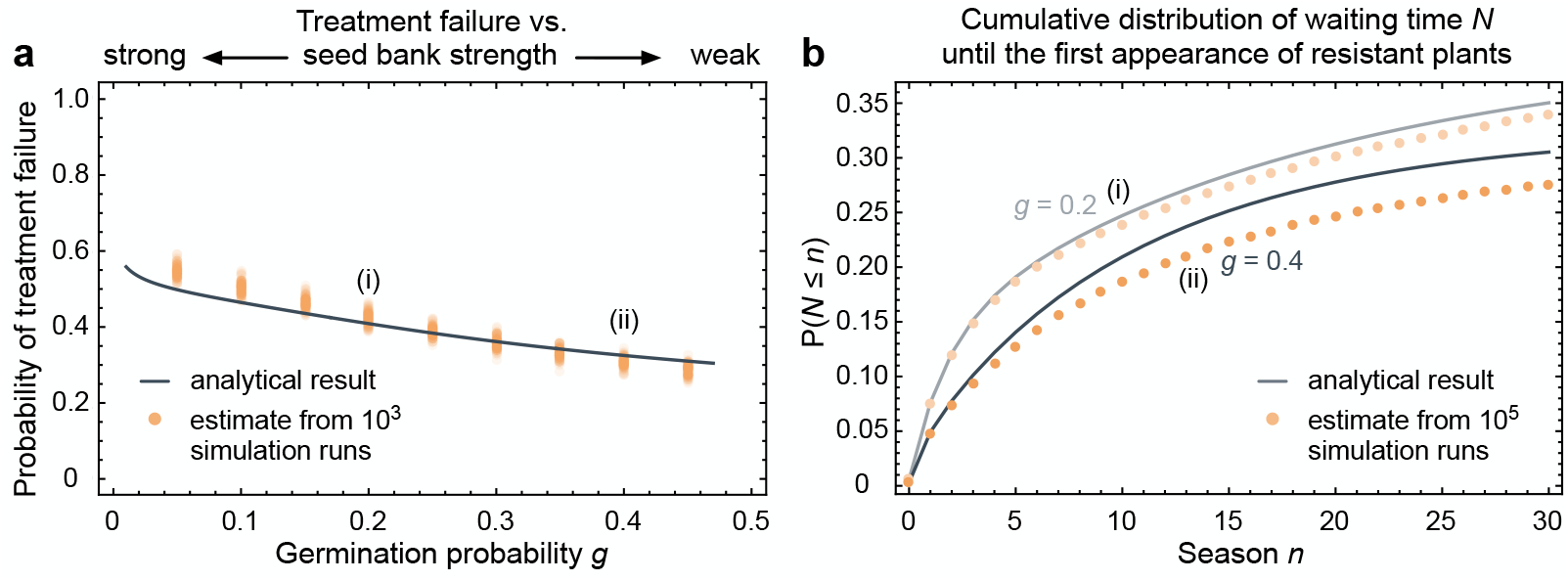
Impact of the seed bank on herbicide resistance adaptation. We show the germination probability’s effect, determining the seed bank’s strength, on control failure. The seed bank mortality was adapted to maintain the total expected number of sexually generated offspring of a plant constant. We simulated a previous herbicide treatment followed by ten years without any control measures and used the result as the start population composition. **a, Variation of the treatment failure probability with the germination probability**. The grey line shows the probability of treatment failure derived analytically from Equation (9). The closed orange circles are estimates from 10^3^ simulation runs with 10^2^ replicates each. **b, Distribution of waiting time until the first resistant plant appears in weed populations with varyingly strong seed banks**. The cumulative waiting time distribution was derived analytically from (15), shown as grey lines. The closed orange circles are estimates from 10^5^ simulation runs. The dark grey line and closed orange circles show the cumulative distribution of waiting time under a weaker seed bank with a germination probability of *g* = 0.4 (ii), and the light grey line and closed light orange circles correspond to the waiting in populations that have a stronger seed bank with a germination probability of *g* = 0.2 (i). The corresponding treatment failure probabilities in (a) are marked accordingly. The figures use the default parameter set as defined in ***Table S1*** except when specified otherwise.

#### Considering self-pollination

Self-pollination increases the level of homozygosity in a population and reduces target-site resistance adaptation from standing variation (as in Subsection Source of adaptive variation). In the early seasons of treatment, resistant plants are more likely to appear in cross-pollinating than in self-pollinating populations (***Figure 6*** a and ***Figure S2*** a). This is caused by the reduced frequency of resistant individuals in the standing variation of self-pollinating weed populations. Resistance alleles cluster in homozygotes, decreasing the number of individuals resistant towards the herbicide unless the mutation is highly recessive. A recessive resistance cost further decreases the number of offspring a homozygous resistant plant generates compared to a heterozygote, diminishing standing variation for target site resistance. In later seasons, where adaptation likely results from *de novo* mutations, no notable effect of the mating system exists on the waiting time distributions for resistant plants (***Figure 6*** a and ***Figure S2*** a). Under cross-pollination, it is unlikely that the first mutant plant to establish is homozygous, while for self-pollinating populations, there might be resistant homozygotes in the standing variation able to rescue the population (***Figure 6*** b). However, mutations arising *de novo* usually occur heterozygously. Resistant homozygotes generally only appear as the first resistant plant on a field in the first seasons of treatment; otherwise, they emerge as offspring of established heterozygotes.

**Figure 6.**
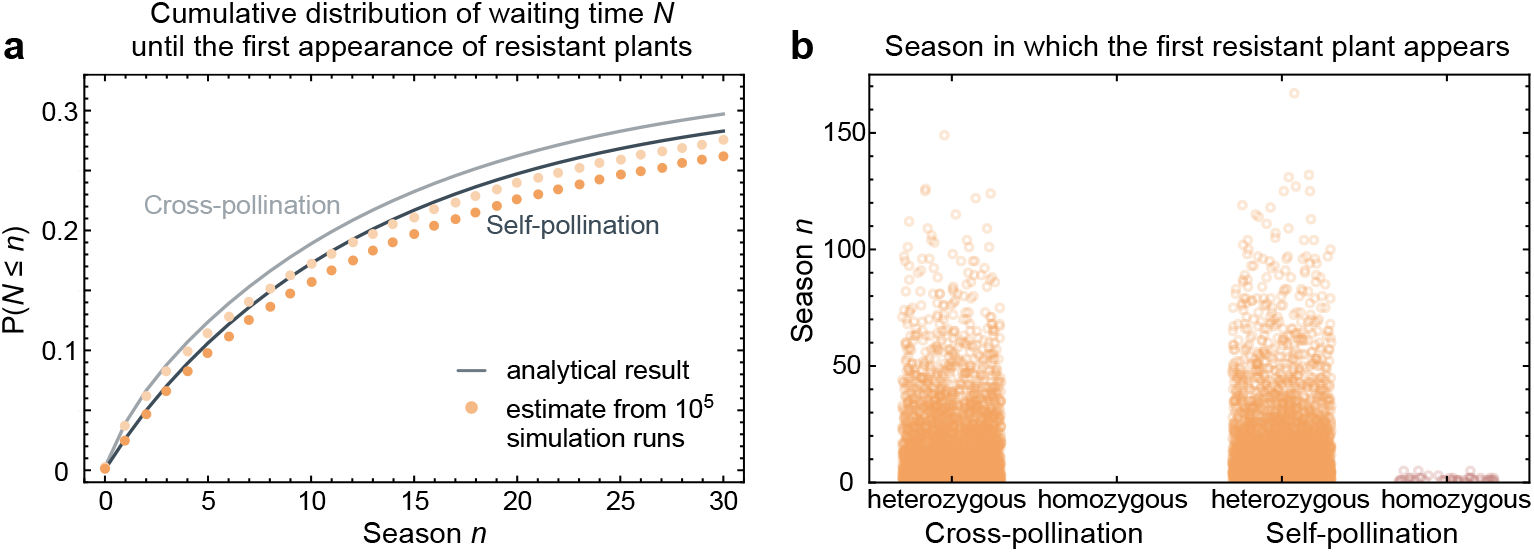
Impact of self-pollination on the timing of resistance evolution. We show the time distribution until the establishment of resistant plants on the field under cross- and self-pollination. **a, Distribution of waiting time until the first resistant plant appears**. The cumulative waiting time distribution was derived analytically from (15), shown as grey lines. The closed orange circles are estimates from 10^5^ simulation runs. The light grey line and closed light orange circles depict the cumulative waiting time distribution in a cross-pollinated population, and the line in dark grey and the closed orange circles correspond to the waiting time in a purely self-pollinated population. **b, Time of the first appearance of a resistant plant**. On the left, we show simulation results for purely cross-pollinating populations and on the right for weed populations that self-pollinate, with 10^4^ replicates each. The open orange circles represent heterozygous resistant plants, and the open red circles show when the first resistant plant surviving on the field is homozygous. The figures use the default parameter set as defined in ***Table S1*** except when specified otherwise.

## Discussion

Various factors influence the primary source of adaptive variation. Using a haploid model focused on adaptation via a single beneficial allele, Orr and Unckless (2014) demonstrated that the probability of adaptation from *de novo* mutations versus standing variation is influenced by the frequency of the beneficial allele and mutation rate and also by the rate of population decline. Faster population declines enhance the role of standing variants, underscoring the potential significance of standing genetic variation for rapid adaptation following abrupt environmental shifts. Hawkins et al. (2019) emphasise the role of the selection coefficient for the source of adaptation, arguing that, in the context of herbicide resistance, mechanisms conferring high levels of resistance, like target-site resistance, can emerge *de novo*. Genomic studies in blackgrass and waterhemp reveal different primary sources of target-site resistance in these two annual weeds (Kreiner et al., 2022; Kersten et al., 2023). We find that fitness costs associated with resistance, the type of pollination, and the weed’s fecundity and seed bank size are other key factors determining the relevance of *de novo* mutations and standing genetic variation for the evolution of herbicide resistance. Thus, we stress the necessity of ecological and evolutionary details in the assessment of strategies for resistance management. The high fecundity of Johnsongrass and the high level of resistance conferred by target-site mutation seem to fuel adaptation from *de novo* mutations. Simultaneously, resistance-related fitness costs and the prevalence of self-pollination reduce the frequency of pre-existing resistant variants.

The probability of a weed population successfully adapting to herbicide treatment is determined by the absolute number of the resistance allele, not its relative frequency. A larger seed bank, potentially containing more resistant seeds, boosts the probability of adaptation from pre-existing variation. The mutational influx increases with increasing fecundity of a weed, amplifying resistance evolution through *de novo* mutations. This accounts for the strong positive effect of seed production on the risk of resistance described by Neve (2008). Resistance costs depleting the standing variation for target-site resistance reduce adaptation from pre-existing variation substantially (see Lauenroth and Gokhale (2023) for a detailed analysis of the standing variation for target-site resistance concerning the associated fitness cost). This result aligns with the theoretical predictions of Neve (2008) for the total herbicide resistance probability. However, not all mutations endowing target-site resistance are known to involve a fitness cost (Menchari et al., 2008). Our finding that self-pollination diminishes the adaptation of target-site resistance from standing genetic variation in a perennial weed aligns with predictions from a fundamental single-locus model by Glémin and Ronfort (2013). However, the substantial contribution of new mutations might lead to similar probabilities of treatment failure for different pollination types. Neve (2008) likewise predict little difference in the probability of herbicide resistance in cross- and self-pollinated populations when resistance is incompletely dominant, as in our study. Despite this, self-pollinating populations, especially where heterozygotes have low fitness, may have a higher chance of mutant allele establishment (Uecker, 2017). This suggests that under certain conditions, the effective generation of mutant homozygotes in selfing populations, increasing the efficacy of selection, could enhance the likelihood of successful adaptation. Apart from the pollination type, the reproduction type is another detail of a weed’s reproductive system that influences adaptation.

Considering a quantitative trait with gametic and somatic mutations, Orive et al. (2017) demonstrated that clonal reproduction increases population survival after a sudden environmental shift if the standing variation allows for persistence. However, for a slightly beneficial allele with unfit heterozygotes, Uecker (2017) showed that whether asexual reproduction is advantageous depends on its dominance. For recessive target-site mutations, sexual reproduction generating fit homozygotes is vital. Our model captures mutations occurring during sexual reproduction but not somatic mutations. Genome-wide mutation rates (per nucleotide per generation) observed in a vegetatively propagated grass species are similar to those in germ cells (Caetano-Anollés, 1999). Thus, our model underestimates the number of *de novo* resistance mutations that arise and, with that, the probability of control failure, especially in populations with high proportions of asexual reproduction. Together with the incomplete dominance of resistance, this might be why we find no direct positive effect of clonal reproduction based on the resistance heredity. Rhizomatous Johnsongrass is, however, less susceptible towards herbicides (Beasley, 1970; Vidrine, 1989; Tuesca et al., 1999), increasing persistence and buying, similar to a seed bank, time for the weed population to adapt. The improved persistence under herbicide treatment is, besides the resistance heredity, a second mechanism by which vegetative propagation functions as a strategy to overcome control in weeds.

Seed banks preserve genetic information of forgone selection for resistance and prolong the period in which a weed can adapt towards a herbicide, emphasising the complications seed banks impose on managing herbicide resistance. A previous study showed that a seed bank substantially delays the extinction of a herbicide-treated weed population and increases the risk of resistance (Lauenroth and Gokhale, 2023). Further, a seed bank buffers the genetic effects of selection, slowing resistance evolution if it occurs. Under a constant seed bank mortality, an increase in the germination probability of seeds increases the risk of resistant types in the seed bank establishing as resistant plants on the field (Neve, 2008). We compensated the increase in germination by the seed bank mortality so that it did not result in a rise in germinated seeds and simulated a forgone treatment with the herbicide. Under these conditions, lower germination helps preserve resistant types in the seed bank, allowing the weed population to adapt readily to subsequent herbicide applications and increasing the risk of treatment failure.

Our model ignores density dependence or resource competition between the plants. This simplifying assumption is reasonable for low initial plant densities but might systematically overestimate treatment failure probabilities if start densities are high (***Figure S3***). Agronomically, it makes sense to manage even minor weed infestations. In general, population density affects the growth of a mutant lineage, determining its establishment probability (Wilson, Pennings, and Petrov, 2017). However, the herbicide drastically reduces the wild-type weed density from the first application onwards, and competition from the surviving susceptible plants against resistant mutants is neglectable. Precisely, the majority of weeds sensitive towards the herbicide are killed by its application when still in an early growth stage (Dogan and Boz, 2005; Johnson and Norsworthy, 2014), such that the population gets reduced before competition from maturing susceptible plants becomes substantial (Williams and Ingber, 1977; Schwinning et al., 2017). Further, branching processes that survive grow to infinity. Though biologically unrealistic, this is no limitation to the results since resistant subpopulations that reached a sufficient size ensure population survival almost certainly (Harris, 1963, pp. 10-11). Therefore, competition between plants of one or more resistant lineages does not influence the results. Density effects in seed production reduce the generation of mutants in the first seasons of herbicide treatment, lowering the probability of resistant plants first appearing in these seasons slightly compared to analytical results from Equation (14) (***Figure S2***). Estimates from simulations implementing density-dependent reproduction match well with the probability of treatment failure derived analytically from Equation (9) for an initial weed density of one plant per square metre (***Figure 4*** a and ***Figure 5*** a). However, for higher weed densities in season 0, the analytical results systematically deviate from the density-dependent simulations, overestimating the probability of treatment failure (***Figure S3*** b). Since, in our model, the start population is untreated, population density affects the number of offspring generated by these initial plants, determining the rescue probability.

Most mathematical models of resistance evolution implement a single biallelic locus in a haploid or diploid population (but see Santer and Uecker (2020) Holmes et al. (2022) and Garoña et al. (2023)). The results presented in this manuscript are for a single locus in a diploid genome. Though we did not show it here, our flexible model can explicitly capture several loci with epistatic interactions and any ploidy level. The higher number of genotypes results in a larger type space of the branching process, increasing the computational complexity. A broad mutational target size increases the probability of treatment failure. Orr and Unckless (2008) observed an effect similar to an increase in population size when there are no epistatic interactions. Furthermore, our model can describe quantitative genetic scenarios often observed for non-target-site resistance, though the type space will become rather large. It would be interesting to study the differences between monogenetic and polygenetic resistance mechanisms and the factors shaping their role in the evolution of herbicide resistance.

Mathematical models of evolutionary rescue, e.g. Orr and Unckless (2008), Orr and Unckless (2014), and Tomasini and Peischl (2020), commonly use Haldane’s (1927) classic approximation of the probability that a unique beneficial mutation escapes stochastic loss in order to obtain analytical results. It assumes the mutant’s absolute fitness to be greater than but close to one. The evolution of resistances in herbicide-treated weed populations is an example of rapid adaptation under strong selection (Kreiner, Stinchcombe, and Wright, 2017). Target-site resistance typically endows a high fitness, not captured by Haldane’s (1927) approximation. Our approach allows for mutants with absolute fitness considerably larger than one, with the drawback that we do not obtain analytical solutions for all considered probabilities. The probabilities calculated in Paragraph Waiting times are analytically exact results, while the extinction and rescue probabilities presented in Paragraph Probability of treatment failure are numerical results. The *m*-dimensional equation system (7) is solved numerically, using Newton methods, to derive the type-specific extinction probabilities ***q*** (FindRoot in Mathematica version 13.1).

We demonstrated how concepts of rescue theory help to understand the factors determining the probability and timing of treatment failure resulting from the establishment of herbicide-resistant weeds. The methods can be employed to identify weedy traits that favour adaptation from *de novo* mutations or standing genetic variation. Sustainable weed management strategies comprise diverse control measures, including crop and herbicide rotations. Studying the evolution of resistance and risk of treatment failure in these fluctuating environments will be of added practical relevance. In addition to new mutations or pre-existing variation, herbicide resistance might also result from gene flow from resistant neighbouring fields. As herbicide susceptibility is a shared resource among farmers, a metapopulation model allowing for pollen, seed and rhizome migration between fields could be an exciting extension of our approach.

## Acknowledgment

We thank Hildegard Uecker, Gaurav Athreya and Demetris Taliadoros, and the research group Theoretical Models in Eco-evolutionary Dynamics for their valuable comments on the manuscript and the research group *Theoretical Models in Eco-evolutionary Dynamics* for helpful discussions. This work is dedicated to the memory of Karsten Keller, an exceptional teacher and mentor known for his kindness and wholehearted support for his students. Although Karsten could no longer contribute to this work directly, his lectures and guidance over the years built its foundation. The preprint was created using the LaPreprint template (https://github.com/roaldarbol/lapreprint) by Mikkel Roald-Arbøl .

## Author contributions

Both authors conceived the study, developed the model, analysed the results and wrote the manuscript. D.L. formulated the mathematical model, the algorithm and ran the simulations.

## Supplementary Information

### Supplementary figures

**Figure S1.**
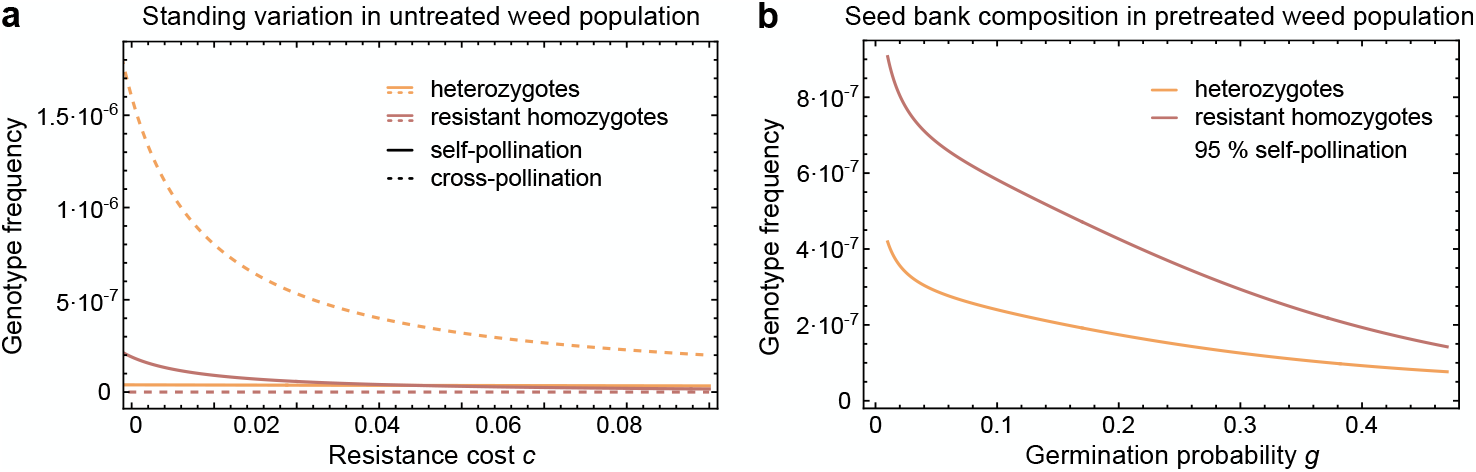
Standing genetic variation for target-site resistance. We used a deterministic density-dependent model (Lauenroth and Gokhale, 2023) to estimate the standing variation for target-site resistance. The relative frequency of heterozygotes is displayed in orange and the relative frequency of resistant homozygotes is shown in red. **a, Standing variation in weed populations never exposed to the herbicide before**. We simulated over 1000 years without management interventions to achieve mutation-selection balance. The equilibrium frequencies are shown for fully self-pollinating populations as solid lines and for cross-pollinating populations as dashed lines. **b, Seed bank composition of a weed population previously treated with the herbicide**. We simulated one year with herbicide application followed by ten years without any control measures, reporting the final seed bank composition. The figures use the default parameter set as defined in Table S1 except when specified otherwise.

**Figure S2.**
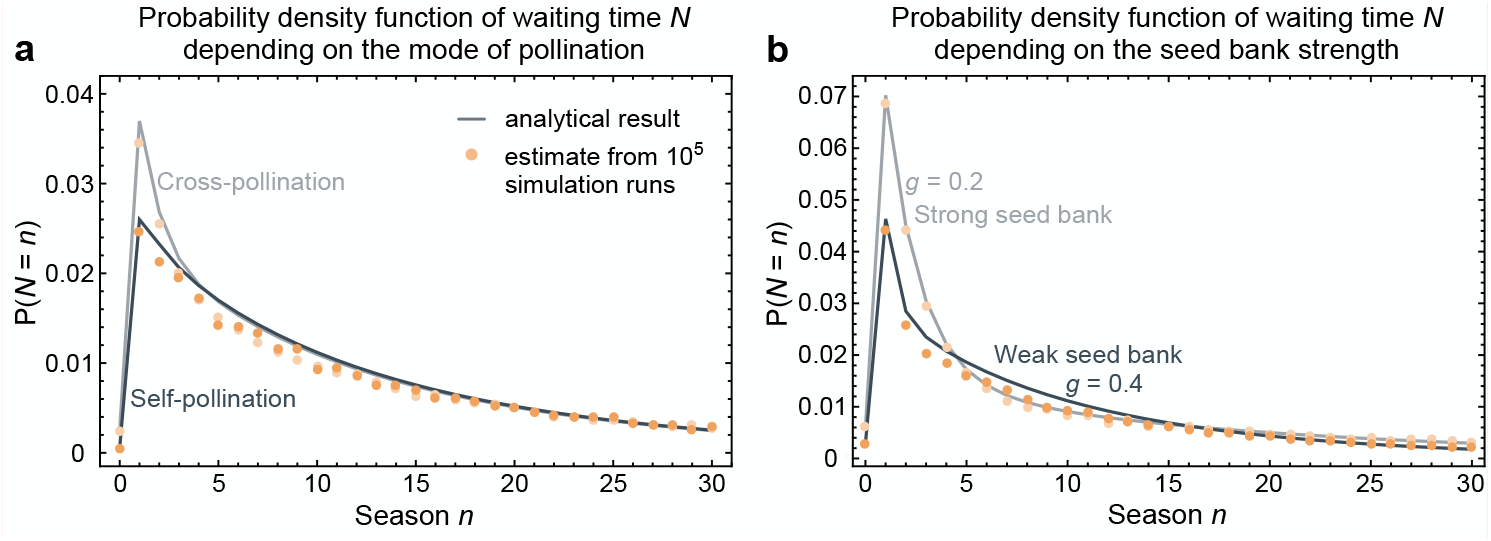
Waiting time until the first resistant plant appears. The probability mass functions of the time *N* until the first appearance of a resistant plant were derived analytically from Equation (15), shown as grey lines. The closed orange circles are estimates from 10^5^ simulation runs. **a, Waiting times in cross- and self-pollinated weed populations**. The waiting time distribution is shown for cross-pollinated populations as a light grey line with closed light orange circles and for self-pollinated populations as a dark grey line with closed orange circles. **b, Waiting times in weed populations with varyingly strong seed banks**. We simulated a previous herbicide treatment followed by ten years without any control measures and used the results as the start population compositions. The dark grey line with closed orange circles shows the probability mass function of waiting time under a weaker seed bank with a germination probability of *g* = 0.4, and the light grey line with closed light orange circles corresponds to the waiting in populations that have a stronger seed bank with a germination probability of *g* = 0.2. The seed bank mortality was adapted to maintain the total expected number of sexually generated offspring, which one plant has, constant. The figures use the default parameter set as defined in Table S1 except when specified otherwise.

**Figure S3.**
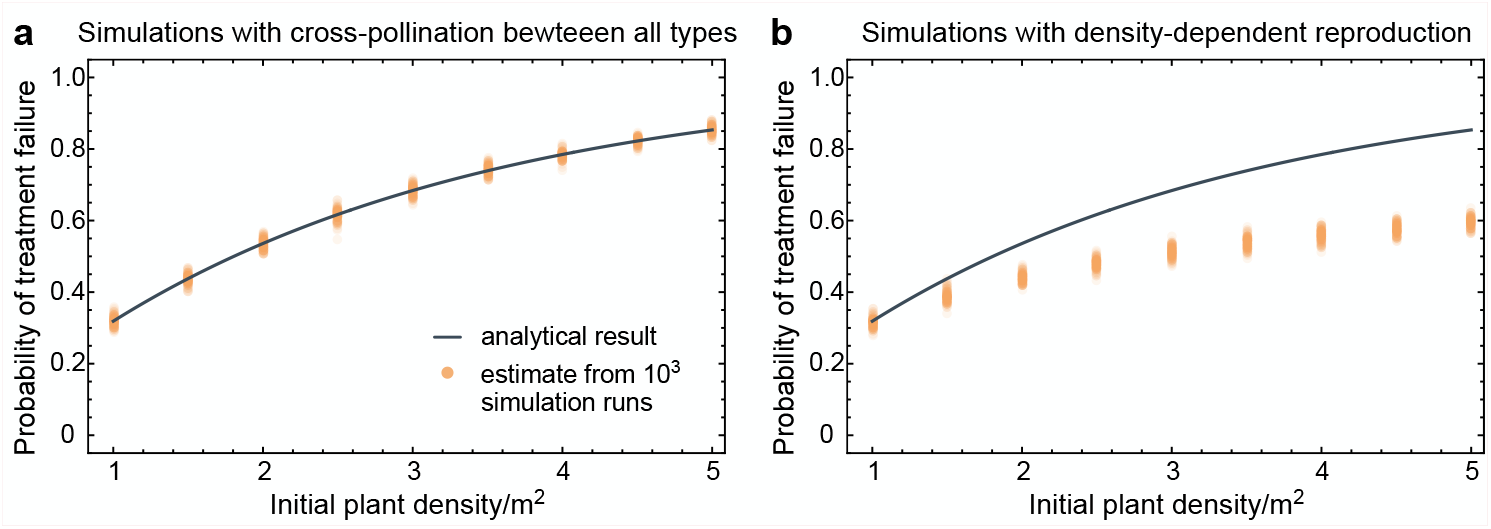
Impact of density-dependence and mating between resistant plants on the probability of treatment failure. We display the variation of the rescue probability with the initial population density. The grey lines show the probability of treatment failure derived analytically from Equation (9), not considering any density effects on reproduction and matings between resistant plants. The closed orange circles are estimates from 10^3^ simulation runs with 10^2^ replicates each. The initial seed density was 80 times the initial plant density. **a, Impact of cross-pollination between resistant plant types on the probability of treatment failure**. We show estimates from simulations comprising cross-pollination between all genotypes. **b, Impact of density-dependent reproduction on the probability of treatment failure**. We show estimates from simulations comprising density-dependent reproduction. The figures use the default parameter set as defined in Table S1 except when specified otherwise.

### Supplementary methods

#### Offspring generating function for *Sorghum halepense*

We illustrate our modelling framework here on the example of *Sorghum halepense* (Johnsongrass). The perennial weed Johnsongrass is a highly competitive invasive species, able to reproduce sexually via seeds and asexually through rhizomes (Warwick and Black, 1983; Peerzada et al., 2023; Schwinning et al., 2017). The plants are dormant throughout the winter, overwintering as seeds and rhizomes in the ground (Warwick and Black, 1983). At the beginning of a growing season, seeds germinate to produce seedlings, and buds on the rhizomes sprout to generate shoots (Peerzada et al., 2023). Mature plants produce new rhizomes and fresh seeds (Warwick and Black, 1983; Peerzada et al., 2023). We illustrate the life cycle of Johnsongrass in ***Figure 1*** a.

The numbers of rhizome buds and seeds produced by a plant of genotype *i* are assumed to be Poisson distributed with mean *b* and (1 − *c*_*i*_) *f*, respectively. Seeds might die over the winter with a given probability *d*_*S*_. In contrast to Lauenroth and Gokhale (2023), here we assume that buds on a rhizome survive the winter independently of each other with a probability (1 − *d*_*Z*_). Therefore, the numbers of surviving rhizome buds and seeds produced by a plant of genotype *i* follow again Poisson distributions with means (1 − *d*_*Z*_) *b* and (1 − *d*_*S*_) (1 − *c*_*i*_) *f* (Acharya et al., 2014). The seeds germinate in the subsequent spring independently of each other to produce seedlings with probability *g* and similarly shoots sprout from the buds on a rhizome with probability *g*_*Z*_. Non-germinated seeds stay in the seed bank, unaffected by potential herbicide application. Seedlings and shoots can die due to herbicide application before reaching reproductive age at given efficiencies *h*_*L*_ and *h*_*T*_, respectively. We illustrate this representation of Johnsongrass’ life cycle in our model in ***Figure 1*** b. The numbers of plants reaching the reproductive stage that originated from a single plant of genotype *i* through asexual propagation and sexual reproduction are consequently Poisson distributed with means *b* (1 − *d*_*Z*_) *g*_*Z*_ (1 − *h*_*T j*_) and (1 − *c*_*i*_) *f* (1 − *d*_*S*_) *g* (1 − *h*_*Lj*_) and independent of each other (Acharya et al., 2014). As the herbicide efficiency might differ between different genotypes, the type-specific efficiencies on seedlings and shoots are denoted by *h*_*Lj*_ and *h*_*T j*_ .

Likewise, the number of seeds in the next season’s seed bank generated by a type *i* plant is independent of its plant offspring and follows a Poisson distribution with mean (1−*c*_*i*_) *f* (1−*d*_*S*_) (1−*g*). While we assume plants only produce rhizome buds of their genotype, seeds might have a different genotype following simple Mendelian inheritance and could include mutations. Let ***M***(*j*), *j* ∈ {1, 2, …, *m*}, be the set of inheritance matrices, where the entries *M*_*ik*_(*j*) give the proportions of type *j* seeds produced by a plant of genotype *i* pollinated by type *k* pollen. Plants self-pollinate at a proportion *p*_self_, corresponding to the pollen-producing plant being of the same genotype as the mother plant (*k* = *i*). We assume that pollen is abundant enough that pollination results in complete ovule fertilisation, neglecting a selective advantage of self-pollination here.

With interest in the establishment of rare alleles, we assume that the sensitive homozygous type, referred to as genotype 1, is abundant, while the types carrying the resistance allele are rare. Therefore, we assume resistant plants do not cross-pollinate with other plants of these types. For type 1 plants, we consider cross-pollination by plants of all possible genotypes. The fraction of sensitive plants pollinated by pollen from a resistant plant type *i ≠* 1 in a season *n* is 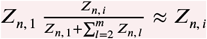 and the fraction cross-pollinated by pollen from sensitive plants is 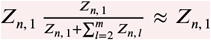. It is, hence, natural to count all offspring from matings between type 1 and rare type plants as the rare type’s offspring. Since the inheritance does not depend on which mating partner contributes the male and which the female gametes, this altered offspring assignment does not constitute a deviation from Mendelian inheritance. Hence, the numbers of genotype *j* plants and dormant seeds produced through sexual reproduction by a resistant plant of genotype *i ≠* 1 are mutually independent and Poisson distributed with mean *f* ((1 − *c*_*i*_) (*M*_*ii*_(*j*) *p*_self_ + *M*_*i*1_(*j*) (1 − *p*_self_)) + (1 − *c*_1_) *M*_1*i*_(*j*) (1 − *p*_self_)) (1 − *d*_*S*_) *g* (1 −*h*_*Lj*_) and *f* ((1 −*c*_*i*_) (*M*_*ii*_(*j*) *p*_self_ +*M*_*i*1_(*j*) (1 −*p*_self_)) +(1 −*c*_1_) *M*_1*i*_(*j*) (1 −*p*_self_)) (1 −*d*_*S*_) (1 −*g*), respectively. Since we consider simple Mendelian inheritance, ***M***(*j*) is symmetric and the respective means reduce to *f* (*M*_*ii*_(*j*) (1 − *c*_*i*_) *p*_self_ +*M*_*i*1_(*j*) (2 − *c*_1_ − *c*_*i*_) (1 −*p*_self_)) (1 −*d*_*S*_) *g* (1 −*h*_*Lj*_) and *f* (*M*_*ii*_(*j*) (1−*c*_*i*_) *p*_self_ +*M*_*i*1_(*j*) (2−*c*_1_ −*c*_*i*_) (1−*p*_self_)) (1−*d*_*S*_) (1−*g*). The numbers of type *j* plants and dormant seeds a type 1 plant has as sexually generated offspring in the next generation are independent and follow Poisson distributions with means *M*_11_(*j*) *f* (1 −*d*_*S*_) *g* (1 −*h*_*Lj*_) and *M*_11_(*j*) *f* (1 − *d*_*S*_) (1 − *g*).

These approximations of the cross-pollination allow us to capture sexual reproduction in the asexual Galton-Watson process and to obtain analytical and numerical results. Since the extinction probabilities of processes starting with a resistant plant type are very close to zero, the survival of a weed population under herbicide control depends almost exclusively on the establishment of resistant plants (see Section Results). The weed survival probability is not notably affected by cross-pollination between resistant types (***Figure S3*** a). For the sexual multitype Galton-Watson process, all theoretical results were derived exclusively for superadditive mating functions (Hull, 1998; Fritsch, Villemonais, and Zalduendo, 2022). However, random mating in a population of several types results in a non-superadditive mating function, not allowing any analysis of the process but solely simulations.

Seeds that did not readily germinate in the first spring after their production die before the following spring with probability *d*_*B*_. Those seeds that stayed in the seed bank germinate in spring with probability *g*. Accordingly, a seed in the seed bank that is of genotype *i* produces one type *i* plant in the next season with the probability (1−*d*_*B*_) *g* (1−*h*_*Li*_) and by staying dormant one seed of genotype *i*—itself—with probability (1 − *d*_*B*_) (1 − *g*). (*d*_*B*_ + (1 − *d*_*B*_) *g h*_*Li*_) gives the probability of a seed disappearing from the seed bank without having any offspring in the next generation.

Knowing the offspring distributions, we can now specify the parameters of the offspring-generating functions for Johnsongrass. For

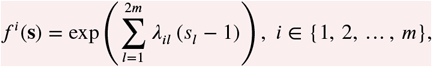

the generating functions of the plant’s offspring,

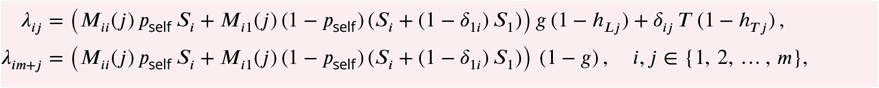

are the parameters of the respective Poisson distributions with *T* = *b* (1 − *d*_*Z*_) *g*_*Z*_ and *S*_*i*_ = (1 − *c*_*i*_) *f* (1 − *d*_*S*_). For the offspring-generating functions of seeds,

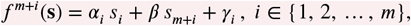

the parameters are given by *α*_*i*_ = (1 − *d*_*B*_) *g* (1 − *h*_*Li*_), *γ*_*i*_ = *d*_*B*_ + (1 − *d*_*B*_) *g h*_*Li*_, *i* ∈ {1, …, *m*}, and *β* = (1 − *d*_*B*_) (1 − *g*).

#### Expected population over time

The offspring generating function ***f*** (***s***) = (*f* ^1^(***s***), *f* ^2^(***s***), …, *f* ^2*m*^(***s***)) can be used to derive the matrix of first-order moments ***A*** with entries

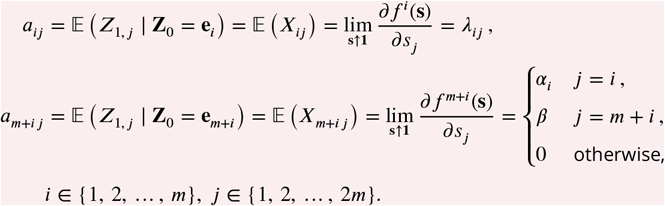

For given genotypes *i, j* ∈ {1, 2, …, *m*} the entries *a*_*ij*_ and *a*_*i m*+*j*_ correspond to the expected numbers of genotype *j* plants and seeds a single plant of genotype *i* leaves in the subsequent season. Likewise, *a*_*m*+*i j*_ and *a*_*m*+*i m*+*j*_ give the expected counts for genotype *j* plants and seeds originating from one genotype *i* seed in the preceding season, where *a*_*ij*_, *a*_*i m*+*j*_ ≤ 1.

Obviously 𝔼 (***Z***_1_ ∣ ***Z***_0_ = ***e***_*i*_) = ***e***_*i*_ ***A*** holds. Using (1) we can more generally express the expectation of ***Z***_*n*+1_, the population in season *n* + 1, conditioned on the population of the previous season ***Z***_*n*_ as

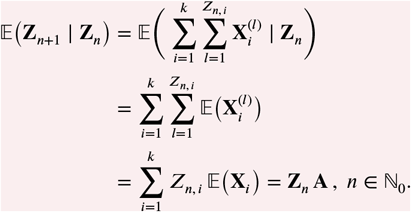

Taking the expectation with respect to ***Z***_*n*_ yields a recurrence formula for the expected population in season *n* + 1 when the initial population is given by ***Z***_0_,

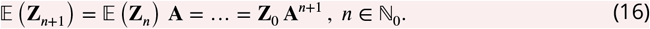

#### Number of mutations

We can derive the number of new mutants in a similar way as Azevedo and Olofsson (2021). Let the random variable *K* ∈ ℕ_0_ describe the total number of distinct resistant plant lineages generated by *de novo* mutation in the lineage of a sensitive plant (type 1). Further, let *v* denote the expectation of *K*. We first need to show that *v* < ∞. Consider a two-type Galton-Watson process 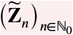 of sensitive plants and seeds, starting with a single plant. The matrix of first-order moments is given by

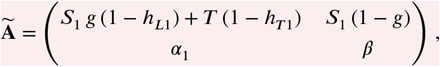

with spectral radius *ρ* < 1 (for the parameter set given in ***Table S1***, *ρ* ≈ 0.935). Let *Y*_*total*_ denote the total number of seeds produced in the lineage of a sensitive plant and *Y*_*n*_ the number of seeds produced in season *n*. Since we only consider mutations occurring in seed production, the total number of new mutants arising is lower than the total number of produced seeds

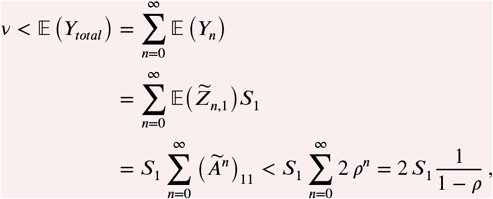

where the third equality follows from Equation (16) and the last inequality is a consequence of the Perron-Frobenius Theorem (Harris, 1963, pp. 37-38).

Let us return to our usual branching process with 2*m* types. Conditioning *v* on the off-spring generated by a sensitive plant, we get the relation

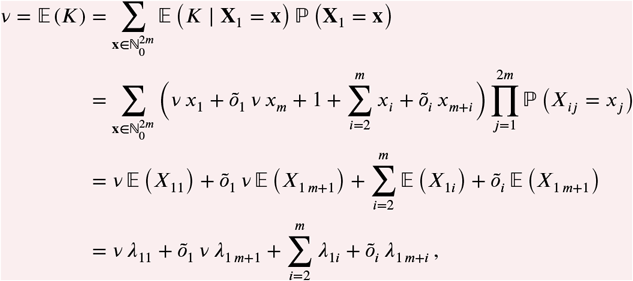

where 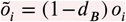 with 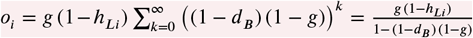 being the proportion of genotype *i* seeds that successfully establish a plant on the field. The third equality follows from the independence of the Poisson distributed offspring, and the fourth equality holds because of the absolute convergence of the series (*v* < ∞). The expected number of new mutants generated in the lineage of a sensitive individual is then given by

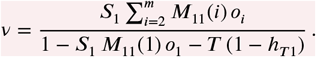

The total number of resistant plants generated by *de novo* mutation in a process starting with Poisson distributed numbers of seeds and plants equals

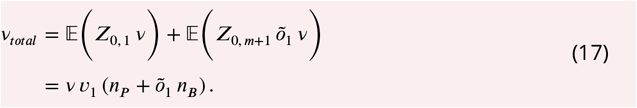

On the other hand, the total contribution of the standing genetic variation to resistant plant lineages appearing on the field is

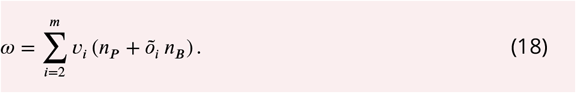

Since 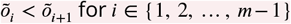, we can derive an upper bound for the ratio of resistant plant lineages emerging in the field that results from *de novo* mutation vs. standing genetic variation

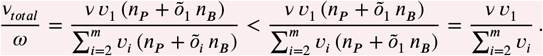

With the parameter set given in ***Table S1***, *de novo* mutations contribute approximately 22 times more resistant plant lineages than the standing genetic variation. However, mutants from the standing variation typically establish earlier, providing an advantage compared to resistant lineages resulting from mutations that appear later in the declining population. Generally, evolutionary rescue from standing variation is faster and requires a smaller population size than rescue from de novo mutations (Orr and Unckless, 2014). While the expected number of resistant lineages resulting from *de novo* mutations *v*_*total*_ is primarily independent of the level of self-pollination (*v*_*total*_ ≈ 0.388), the contribution of the standing variation *ω* is determined by the pollination mode (*ω* ≈ 0.016 in a purely self-pollinating population and *ω* ≈ 0.053 in a fully cross-pollinated population) (***Figure S4***). While new mutations arise unaffected by the source of pollen, self-pollination increases the level of homozygosity in a population, reducing the frequency of standing variants for target-site resistance as resistance alleles cluster in homozygotes. If the mutation is highly recessive, the increased homozygosity, however, yields more resistant individuals compared to outcrossing populations.

**Figure S4.**
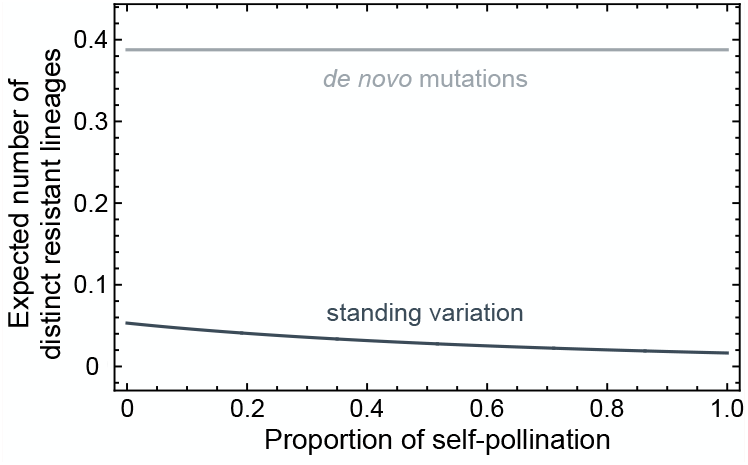
Origin of resistant lineages. We show the effect of self-pollination on the contributions of *de novo* mutations *v*_*total*_ (light grey line) and standing genetic variation *ω* (dark grey line) to the resistant plant lineages appearing on the field. The expected numbers of distinct resistant lineages were derived analytically from Equations (17) and (18). The figures use the default parameter set as defined in Table S1 except when specified otherwise.

#### Computer simulations

We used simulations to analyse the impact of density effects and cross-pollination between resistant plants on the obtained probability and time of herbicide resistance evolution. We implemented our Galton-Watson model with modifications in the sexual reproduction. Our simulations allowed cross-pollination between all possible genotypes and included density-dependent fecundity (as per Eqs. (11) and (12) in Lauenroth and Gokhale (2023)). The plant and seed offspring of type *i* plants followed Poisson distributions with population-dependent means

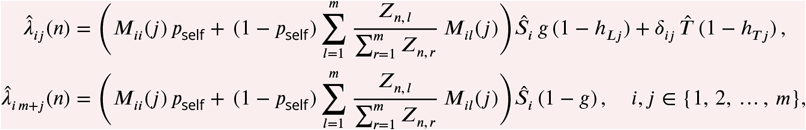

with 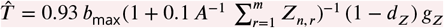 and 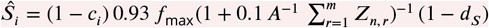 .

### Parameterisation

The results presented in this study are for a single locus in a diploid genome with two alleles—a sensitive allele and an allele conferring herbicide resistance. The sensitive genotype corresponds to genotype 1, the heterozygotes are referred to as genotype 2, and genotype 3 is the homozygous resistant genotype.

#### Standard parameter set

***Table S1*** shows the parameter set of our model with the standard values used for this study. We fixed the parameters displayed in black throughout the whole study. The grey parameters varied in some figures, but take the values indicated here if we do not state them explicitly. We considered incomplete dominance of resistance and the associated fitness cost, such that the type-specific herbicide efficacy and fitness costs are *c*_1_ = 0, *c*_2_ = 0.5 *c, c*_3_ = *c* and *h*_*\**1_ = *h*_*\**_, *h*_*\**2_ = 0.5 *h*_*\**_, *h*_*\**3_ = 0.

#### Inheritance matrices

We suppose simple Mendelian inheritance with mutations in both directions occurring with probability μ. The inheritance matrices in the sexual reproduction of plants are then given by,

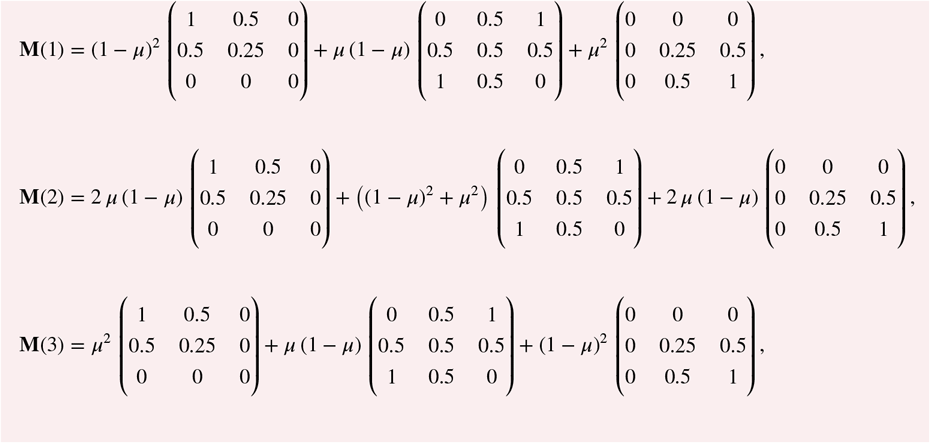

where the entries *M*_*ik*_(*j*) give the proportions of type *j* seeds produced by a plant of genotype *i* pollinated by type *k* pollen.

**Table S1.** Model parameters and their values. Parameters in black are fixed in all calculations and simulations. The parameters displayed in grey are varied in some parts and take the values given here if not stated differently. Detailed explanations and references are given in the Supplementary Information of Lauenroth and Gokhale (2023).

| Model parameter | Symbol | Value |
| --- | --- | --- |
| Sprouting probability | $g_Z$ | 0.2 |
| Germination probability | $g$ | 0.3 |
| Fitness cost on seed production | $c$ | 0.3 |
| Proportion of self-pollination | $p_{\text{self}}$ | 0.95 |
| Mutation rate | $\mu$ | $10^{-8}$ |
| Herbicide efficacy |  |  |
| on seedlings | $h_L$ | 0.998 |
| on shoots | $h_T$ | 0.985 |
| Maximum rhizome bud number produced per plant | $b_{\text{max}}$ | 140 |
| rhizome bud number produced per plant<sup>a</sup> | $b$ | $0.9 b_{\text{max}}$ |
| Maximum viable seed number produced per plant | $f_{\text{max}}$ | 13,000 |
| viable seed number produced per plant<sup>a</sup> | $f$ | $0.9 f_{\text{max}}$ |
| Rhizome winter mortality | $d_Z$ | 0.35 |
| Seed loss |  |  |
| fresh seeds | $d_S$ | 0.94 |
| old seeds in seed bank | $d_B$ | 0.48 |
| Initial density (m<sup>-2</sup>) |  |  |
| seeds |  | 80 |
| plants |  | 1 |
| Field size (m<sup>2</sup>) | $A$ | 10,000 |

<sup>a</sup> In the simulations, the limits of seed and rhizome bud numbers produced per plant are $0.93 f_{\text{max}}$ and $0.93 b_{\text{max}}$ to compensate for the fecundity reduction by $(1 + 0.1 A^{-1} \sum_{i=1}^m Z_{n,i})^{-1}$ apparent even at extremely low densities (Lauenroth and Gokhale, 2023).

